# Benchmarking computational decontamination of ambient RNA

**DOI:** 10.64898/2026.01.13.699237

**Authors:** Cecilie Bøgh Cargnelli, Jakob Vennike Nielsen, Jesper Grud Skat Madsen

## Abstract

Gene expression profiling of single cells using single-cell and single-nucleus RNA sequencing (sxRNA-seq) enables researchers to characterize cellular heterogeneity and unraveling complex biological processes at unprecedented resolution. However, sxRNA-seq faces challenges due to the presence of ambient RNA, extraneous RNA molecules not originating from the cells of interest. Sample preparation is a major source of ambient RNA, where harsh conditions can lead to cell lysis and the release of intracellular RNA. This inescapable inclusion of ambient RNA can cause erroneous results and hinder downstream analyses. To address this issue, various methodologies have been developed to identify, quantify, and remove ambient RNA. Here, we rigorously evaluate 7 state-of-the-art methodologies for ambient RNA removal using simulated datasets, species-mixing experiments of varying complexities, and genotype-mixing experiments. We find that no single method performs the best across all datasets and metrics, but CellBender, DecontX and SoupX generally perform well.

## Introduction

In recent years, single-cell and single-nucleus RNA-sequencing (sxRNA-seq) have revolutionized the field of genomics by enabling the interrogation of gene expression profiles at the resolution of individual cells. This technology enables researchers to unravel cellular heterogeneity, identify rare cell populations, and shed light on dynamic cellular processes. However, despite its transformative potential, sxRNA-seq faces several significant hurdles.

One of hurdles is the presence of ambient RNA. Ambient RNA is defined as extraneous RNA molecules that are present in the experimental environment, but do not originate from the cell of interest. It is likely that especially during sample preparation, where harsh experimental conditions using enzymatic digestion, physical shearing or detergents can lead to unintentional cell lysis or membrane damage, contaminating intra-cellular RNA molecules may be released. Additional sources of contamination include barcode swapping in multiplexed experiments, where molecules from one sample can be incorrectly labelled with a cell barcode from another sample^1^, cross contamination between wells^2,3^ and contamination from the environment, such as media or reagents.

The inadvertent inclusion of ambient RNA in sxRNA-seq data can lead to spurious results, impede accurate cell classification, and confound downstream analyses^4,5^. To tackle the challenge of ambient RNA in sxRNA-seq, datasets can be improved by computational decontamination which aims to identify and quantify ambient RNA, distinguishing it from endogenous RNA, and ultimately remove it to minimize its impact on results. While several decontamination methods have been introduced, no independent benchmark covering all currently available methods across multiple datasets is available to highlight their strengths and limitations. This is important for developers to pinpoint critical areas for further development, and for researchers to select the appropriate tool for the task.

Here, we present a comprehensive and independent benchmarking of 7 computational decontamination methods (CellBender, DecontX, FastCAR, scAR, scCDC, SoupX, and CellClear)^3,6–11^ using both real and synthetic datasets of different complexities, as well as negative control datasets without ambient contamination. We evaluate the ability of each method to remove ambient RNA and preserve endogenous biological signals, and the effect thereof on biological findings. No single method performs the best across all datasets and metrics, yet we recommend using either CellBender, DecontX or SoupX depending on compute environment, data availability and prior expectation.

## Results

### Strategy for benchmarking ambient RNA removal methods

Because ambient RNA contamination is pervasive in single-cell and single-nucleus RNA-sequencing (sxRNA-seq) data, and because it can systematically bias cell-type identification and gene-level inference, a careful benchmarking of ambient RNA removal methods is necessary to understand their practical impact on downstream analyses. Accordingly, we benchmarked the ability of CellBender, DecontX, FastCAR, scAR, scCDC, SoupX, and CellClear^3,6–11^ to remove ambient RNA while preserving endogenous gene expression. Further, we evaluated how ambient RNA removal influences biological interpretation by evaluating clustering, label transfer, and marker gene quality (Figure 1). To define ground truth profiles, we simulated datasets with varying levels of ambient RNA and analyzed strain-mixing and genotype-mixing experiments, leveraging the different biological origin of each read to establish ground truth (Supplemental Table S1).

**Figure 1:**
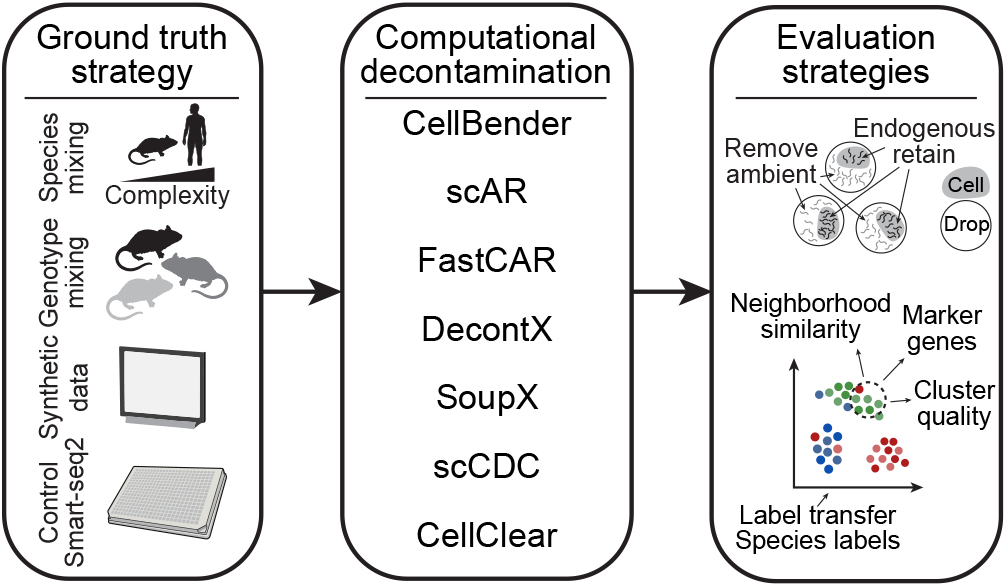
Setup for benchmarking ambient RNA removal methods. Schematic diagram showing the benchmarking workflow. Three species-mixing, genotype-mixing, simulated, and negative control datasets were processed using 7 state-of-the-art ambient RNA removal tools and evaluated in terms of their ability to remove ambient RNA, conserve endogenous gene expression and effect on cell type labelling, clustering, and markers. Based on these, we provide recommendations for different methods depending on which types of data are available.

Following incorporation of computational decontamination methods into standardized scRNA-seq preprocessing workflows, such as nf-core^12^, they are increasingly being applied as a routine background correction step. While this has improved accessibility, it has also lowered the threshold for applying probabilistic background correction without proper prior dataset-specific evaluation, raising the risk of over-correction. To explicitly assess the consequences of applying ambient RNA removal methods without prior evidence of contamination, we also simulated datasets without contamination and included a smart-seq2-based dataset^13^ as negative controls. This allowed us to assess distortion of gene expression and downstream biological interpretation in settings where decontamination is not warranted.

### Ubiquitously expressed messenger RNAs contribute most to ambient RNA

We started out by investigating how variable ambient RNA contamination is between cells in the same dataset and between datasets. To that end, we calculated the ambient load (the fraction of total UMIs derived from an ambient source) in the three species-mixing datasets, as well as the murine strain-mixing dataset. We found that the ambient load is generally higher in the single-nucleus RNA-seq samples (average: 11.80%) than the single-cell RNA-seq samples (average: 7.79%) consistent with previous observations^16^. Moreover, we found that the fraction of ambient RNA per cell after quality control varies significantly within samples (average quartile coefficient of dispersion (QCD) = 0.314), between samples within the same study (average QCD = 0.319), and between studies (average QCD = 0.171) (Figure 2A). Across features, the ambient RNA load (i.e., the fraction of UMIs derived from ambient RNA molecules) is strongly and positively correlated (average R = 0.857) with the average expression level of the same feature in “empty” droplets (Figure 2B), as well as with average expression level across all cells (Supplemental Figure 1A). The ambient RNA load is also non-linearly associated with endogenous expression frequency (Figure 2C), as the 10% most frequently expressed features contribute 82.30% of ambient RNA on average. We observed a similar non-linear association with the fraction of UMIs derived from exons (Supplemental Figure 1B), where the features with 10% highest exonic fractions contribute 24.60% of ambient RNA on average. This suggests that ubiquitously expressed mature cytoplasmic messenger RNAs is the major source of ambient RNA.

**Figure 2:**
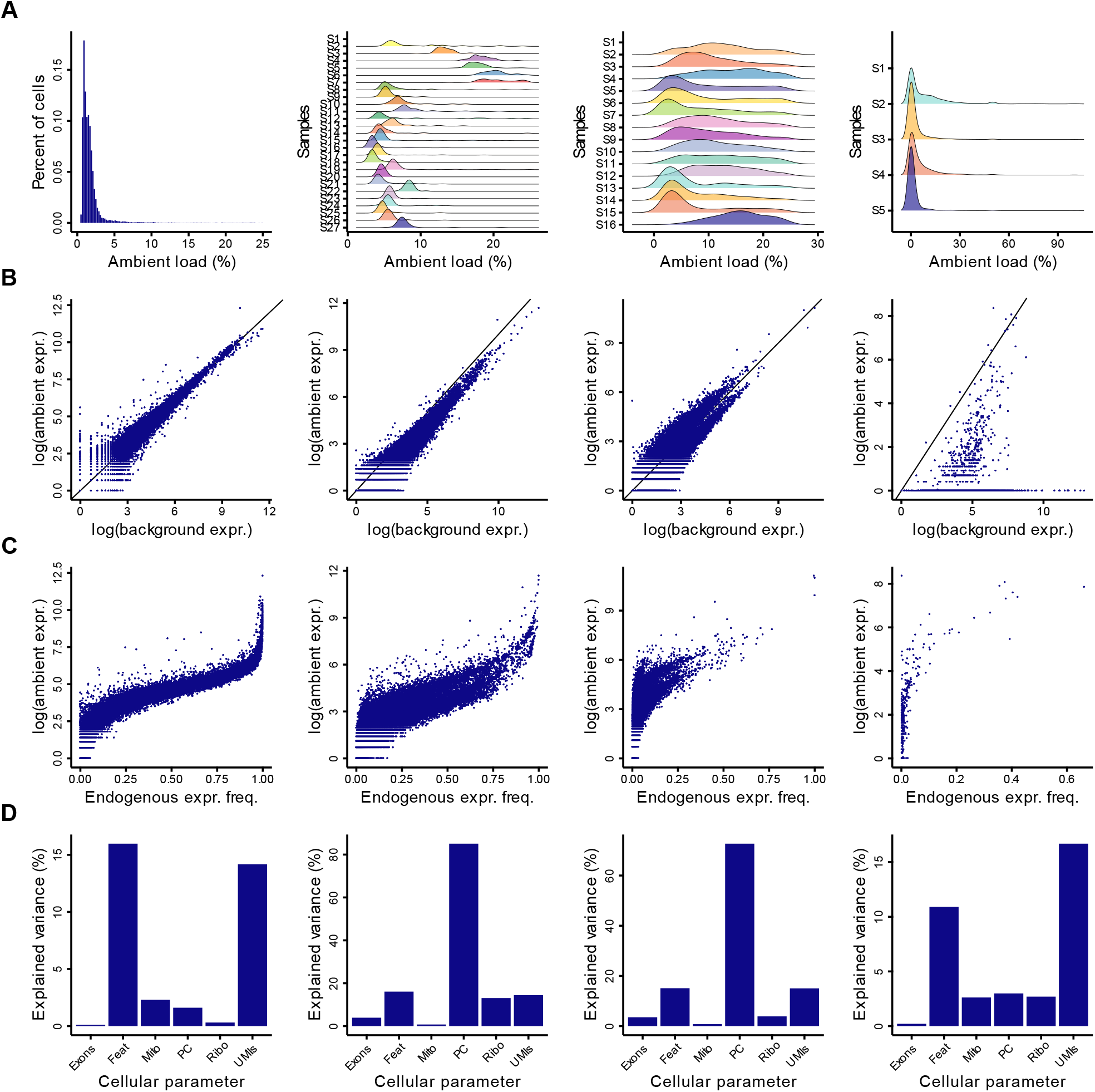
Characteristics of ambient RNA. **A)** Histogram or ridge plots of ambient load (fraction of total UMIs not derived from the cell) for each cell for the indicated datasets and samples.**B)** Scatterplot showing the ambient load on genes and their expression level in empty droplets. **C)** Scatterplot showing ambient load on genes and their endogenous expression frequency. **D)** Barplot showing the percentage of variance in ambient load across cells explained by cellular parameters, including the fraction of UMIs mapped to exons (Exons), mitochondrial genes (Mito), to protein-coding gens (PC), to ribosomal genes (Ribo), as well as the total number of UMIs (UMIs) and features detected (Feat).

Across cells, the ambient load is generally only weakly associated to commonly used cell quality metrics, such as fraction of UMIs derived from exons, or from either mitochondrial or ribosome genes. However, for two datasets, there is a strong association to the fraction of UMIs derived from protein-coding genes, and for all datasets, there is an inverse association with the number of UMIs and the number of unique features (Figure 2D, Supplemental Figure 1C-D). Thus, ambient RNA can be reduced through increasingly stringent quality filtering, but the problem cannot be resolved, as even among the cells with the 10% largest number of UMIs, 2.64% are strongly affected (>10% of UMIs) by ambient RNA.

### Balance of ambient RNA removal and endogenous RNA retainment

To compare different methods for ambient RNA removal, we calculated two scores for each dataset (see Methods), one for removal of ambient RNA and one for retaining endogenous RNA. Both scores range between 0 and 1, where 1 is the best possible performance and 0 is the worst possible performance. In terms of ambient RNA removal, scAR (average = 0.679) and CellBender (average = 0.604) have the best performance (Figure 3A-B, Supplemental Table S2). However, scAR also removes a substantial proportion (average = 0.445) of endogenous RNA signal, especially from lowly expressed genes (Figure 3C). Generally, we observe a positive association between the expression level of genes and ambient RNA removal (i.e., ambient RNA is predominantly removed from highly expressed genes) (Supplemental Figure 2), while we found a negative association with endogenous RNA retainment (i.e., endogenous signal is predominantly removed from lowly expressed genes) (Supplemental Figure 3). In control datasets without ambient RNA, all methods remove approximately the same amount of signal from endogenous RNA, as compared to datasets with ambient RNA.

**Figure 3:**
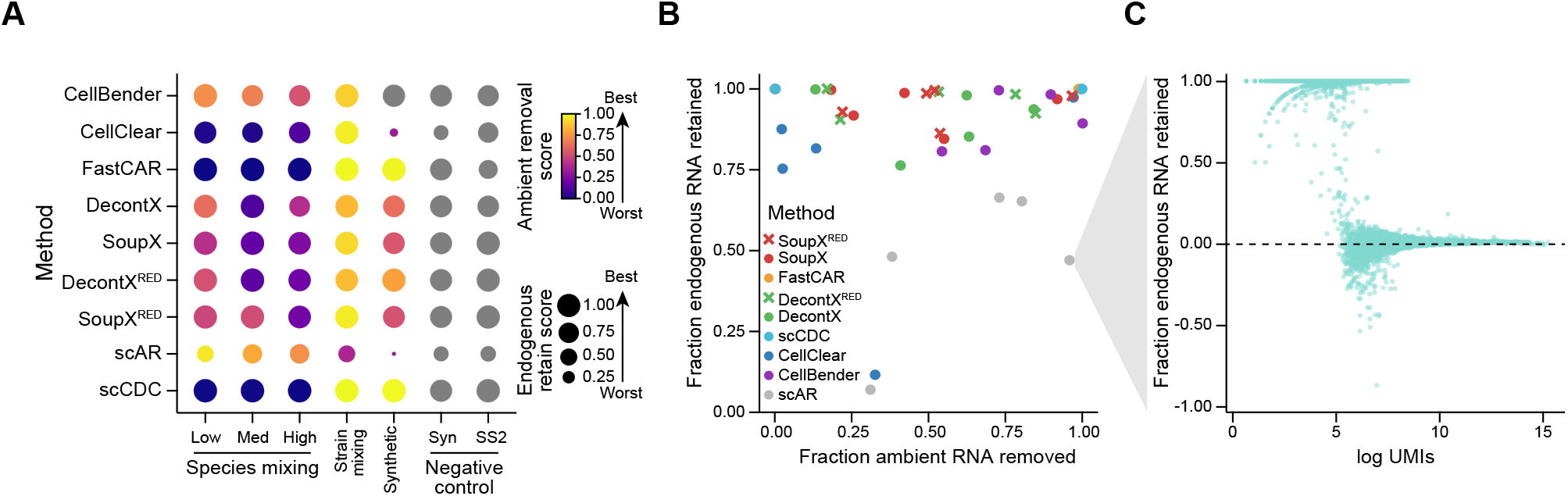
Method performance. **A)** Dotplot of ambient removal scores (shown by color) and endogenous retain scores (shown by dot size) for all seven methods. For SoupX and DecontX, reduced mode (RED) runs were also included that only use filtered matrices across all seven datasets. **B)** Scatterplot showing the fraction of endogenous RNA retained and ambient RNA removed for all datasets. The color of the dot indicates the method. **C)** Scatterplot showing the fraction of endogenous RNA retained and the logged sum of UMIs for each gene after scAR correction in the low complexity species-mixing dataset.

An important distinction between the methods is whether they require access to the full count matrix or can be applied using only the filtered matrix (subset of the full matrix where low quality and empty droplets have been removed), as the full count matrix may not be readily available from public datasets where access to the raw data is restricted. Of the 7 methods, four (SoupX, DecontX, scAR, and scCDC) are applicable to datasets with access to filtered count matrices only, whereof scAR, SoupX and DecontX can be run either using the full count matrix or in a reduced mode using only the filtered matrix. For scAR, the computational resources required to run in full mode were intractable and the analysis did not finish. SoupX in reduced mode removes more ambient RNA compared to full mode, and retains approximately the same amount of endogenous RNA, while DecontX in reduced mode removes slightly less ambient RNA, but retains more endogenous RNA than full mode.

### Effect of decontamination on biological interpretation

To investigate how ambient RNA removal tools impact biological interpretation, we initially evaluated the synthetic sample with the largest ambient RNA load. Only CellBender improves the ability to separate the ground truth cell types using unsupervised clustering relative to using uncorrected counts (Figure 4A). DecontX does not impact cell type recovery, while the remaining methods all perform worse than using uncorrected counts. In terms of marker gene expression, we observed that only DecontX and scAR improves the module scores to approximately the ground truth level (Figure 4B), while most other methods do not significantly improve module scoring above the uncorrected counts. These methods also improved average modules scores in the absence of ambient RNA contamination (Supplemental Figure 4A) indicating a bias towards cell type-specific gene expression.

**Figure 4:**
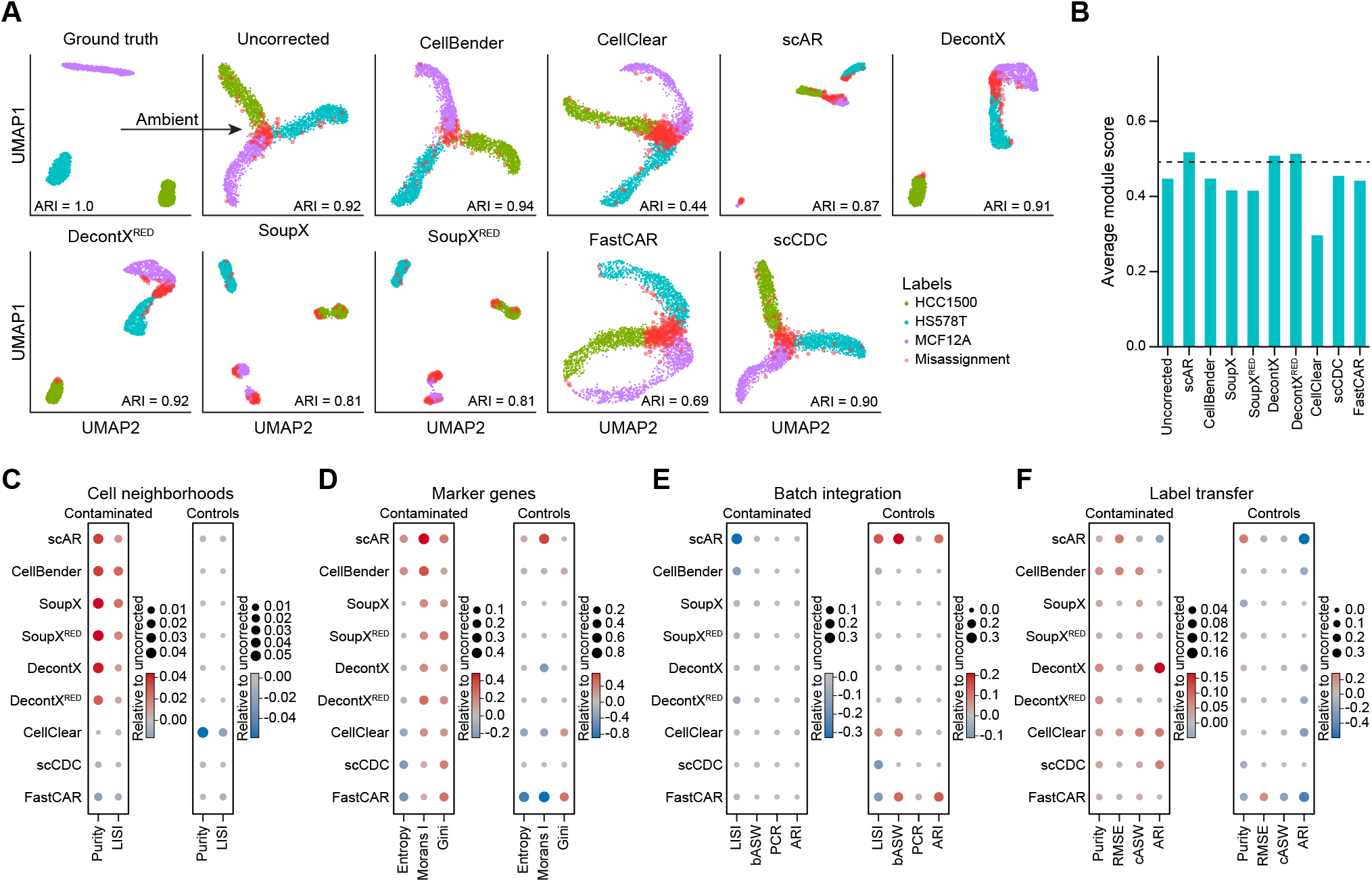
Influence of ambient RNA on biological interpretation. **A)** UMAP embeddings of a synthetic dataset showing the ground truth, as well as the observed data after ambient contamination (uncorrected) and after correction of the observed data for the indicated methods. Each dot was colored in cell type colors if correctly assigned using unsupervised clustering, or in red if misassigned. **B)** Barplot showing average module scores for marker genes defined in the ground truth dataset from A. **C-F)** Dotplots showing the average of the indicated metrics relative to the uncorrected data for contaminated or control datasets across four different tasks: cellular neighborhood similarity (C), marker gene selectivity (D), batch integration (E) and label transfer (F).

To more comprehensively evaluate how computational decontamination methods affect biological interpretation, we evaluated cellular neighborhoods, marker genes, batch integration and label transfer for both contaminated and negative control datasets (Supplemental Table S3-4). For evaluating the quality of cellular neighborhoods, we used species or cell type labels to calculate the purity and local inverse Simpsons index (LISI) for the nearest neighbors of each cell and found that CellBender, scAR, SoupX and DecontX improves the quality of cellular neighborhoods in contaminated datasets and have a negligible impact when applied to negative controls relative to using uncorrected counts (Figure 4C). In terms of marker genes, especially scAR and CellBender leads to more focal expression of marker genes in reduced dimensional space as indicated by improved entropy score (i.e., neighbors have more similar expression patterns), improved Moran’s I (i.e., higher spatial autocorrelation) and Gini coefficient (i.e., markers are expressed by a more selective set of cells) relative to uncorrected counts (Figure 4D). Next, we integrated samples using Harmony^14^ and evaluated integration quality using the average silhouette width across batches (bASW), the batch LISI, principal component regression of batch labels (PCR) and the maximum adjusted Rand index (ARI) between unsupervised clustering and batch labels. We observed that most methods did not significantly improve batch integration relative to using uncorrected counts, but especially scAR led to worse integration, and had a significant impact on integration of negative control datasets without ambient RNA (Figure 4E). Finally, we evaluated the impact of ambient RNA correction on label transfer using Azimuth^15^. We evaluated the purity of transferred labels in local cellular neighborhoods, the average silhouette width (cASW) of transferred labels, the ability to recover them using unsupervised clustering, and the expression similarity between cells assigned to the same cell type label as measured by the root-mean-square-error (RMSE) between each cell and the average profile of all cells within a label. We found that especially, scAR, CellBender, DecontX and CellClear improved label transfer relative to using uncorrected counts, but that scAR had a significant impact on label transfer in datasets without ambient RNA contamination (Figure 4F).

## Discussion

Ambient RNA is pervasive. Across 97.357 cells and nuclei in 49 real samples, we found that 81.56% of cells and nuclei contain some level of ambient RNA. The level varies between cells or nuclei within a sample, between samples within a dataset and between datasets. We found that the ambient load is generally higher in the single-nucleus RNA-seq samples (average: 11.80%) than the single-cell RNA-seq samples (average: 7.79%) consistent with previous observations^16^. Within samples, we did not find any community-standard cell quality parameters that were strongly predictive of the ambient load within an individual cell or nucleus suggesting that ambient RNA load is a stochastic sampling process that is not directly related to the quality of the nuclei or cell. However, the profile of ambient RNA can be accurately estimated using aggregated expression of both empty droplets and non-empty droplets, and it is associated with the expression frequency and exon fraction in non-empty drops.

Across datasets, we found that scCDC and FastCAR primarily removes ambient RNA from highly expressed genes. This skew leads to percentage of ambient RNA being removed, but simultaneously also limits removal of endogenous RNA. We did not observe a positive impact on biological interpretation after applying these methods, which could be due to their expression level bias. CellClear exhibits a similar low ambient RNA removal but has the inverse bias removing ambient RNA primarily from lowly expressed genes. CellClear also significantly alters the endogenous profile, removing approximately as many endogenous RNA counts as ambient RNA counts. DecontX and SoupX rank in the middle in terms of ambient RNA removal and overall, preserves endogenous RNA well. Although SoupX has a slightly larger bias towards being more effective at removing ambient RNA from highly expressed genes than DecontX, they both exhibit a low bias, and they both can improve biological interpretation. Finally, CellBender and scAR are the most effective at removing ambient RNA. Both methods tend to introduce new counts for lowly expressed genes, but unlike CellBender, which generally preserves endogenous RNA and positively impacts biological interpretation, scAR aggressively removes endogenous RNA counts leading to removal of a large percentage of cells in several samples and does not have a strong positive impact on biological interpretation.

In conclusion, we advise the use of either CellBender, DecontX in full mode or SoupX in reduced mode. The choice between the three should be guided by several criteria: expected performance, prior expectation for contamination, usability and scalability (Figure 5, Supplemental Table S5). All methods have adequate documentation. However, only SoupX in reduced mode can be applied to datasets without access to a full count matrix, and CellBender has the highest demand for computational resources, requiring access to a GPU, long processing times, higher peak memory requirements and calls putative cell-containing droplets. The prior expectation for contamination is important, as CellBender and DecontX in full mode are more prone to over-correction in the absence of ambient RNA than SoupX in reduced mode. The prior expectation should be informed experimentally, for example single-nucleus RNA-seq generally have more ambient RNA contamination than single-cell RNA-seq^17^ and potentially computationally, where methods such as AmbiQuant^17^ may be used to infer sample-level contamination. Finally, the expected performance can be inferred from our benchmark, and although no method is universally best, CellBender has the best overall performance, which is driven by ambient RNA removal and retaining endogenous RNA (Figure 5) followed closely by DecontX in full mode, which removes less ambient RNA, but improves biological interpretation more than CellBender and then SoupX in reduced mode. Thus, for a contaminated dataset, we recommend CellBender or DecontX in full mode. Finally, if there is no prior expectation for contamination, or only filtered count matrices are available, we recommend SoupX in reduced mode. However, we do note that applying these methods does not always improve data quality, even in contaminated datasets. In addition to guiding method choice, our benchmark can serve to guide developers to build better computational decontamination methods. To that end, we have made our approaches entirely open source, such that the benchmark can be continuous, adding new tools and updating existing ones as they are evolving, as well as open for parameter optimization.

**Figure 5:**
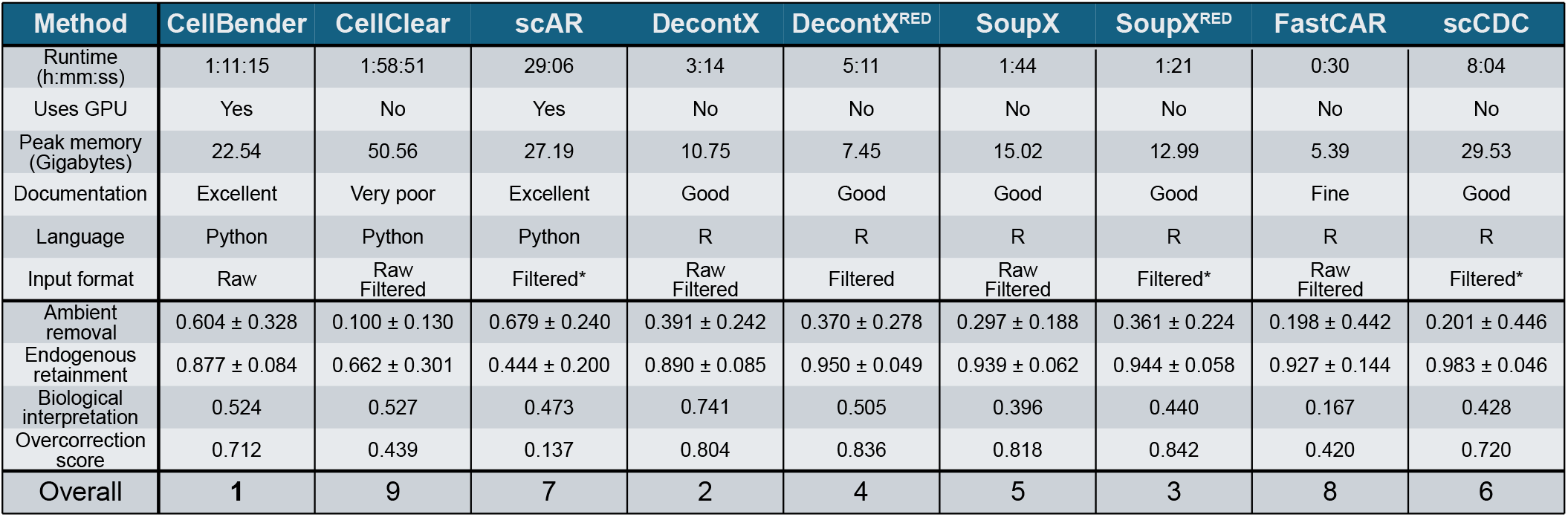
Summary of benchmark. Qualitative and quantitative characteristics of each tested methods summarized across datasets and tasks.

In addition to algorithmic advances, a key limitation in building better decontamination methods is the availability of suitable datasets, where ambient RNA can be quantified in an unbiased manner. Here, we have used multiple complementary approaches for defining ground truth, each with distinct strengths and limitations. We used synthetic data generated by sampling reads from pure cell line expression profiles, mixed in defined ratios. While this provides a well-defined ground truth, the sampling approach might be too simplistic as it does not capture higher-order structures, such as gene-gene covariance, that are present in real data, which may influence decontamination performance. We also used species mixing experiments, where cells or tissues originating from different species are mixed prior to performing the experiment. The data is aligned to a hybrid genome, and ambient RNA is quantified using reads mapping to species X genes within cells assigned to species Y. This provides a robust ground truth if ambitiously aligned reads are excluded. However, the feature spaces of endogenous and contaminating RNA molecules do not overlap, which differs from real datasets and may bias decontamination performance. Finally, we used a strain mixing experiment, where tissues from different strains of mice were mixed prior to the experiment. Here, ambient RNA is identified through strain-specific SNPs detected in cells from another strain. This enables accurate measurement of contamination, but only for a subset of genes, limited by expression levels and genetic variation, which may also introduce bias. Taken together, these complementary strategies help mitigate individual limitations and enable more balanced performance assessment. However, further progress will require new experimental designs in which endogenous and ambient RNA share the same feature space, while contamination can still be measured accurately and without bias across all genes.

## Methods and materials

### Decontamination methods

A total of seven different ambient RNA decontamination methods were included in the benchmark; CellBender^6^ (version 0.3.2), scAR^18^ (version 0.7.0), CellClear^10^ (version 0.0.3), scCDC^11^ (version 1.4), FastCAR^9^ (version 0.1.0), SoupX^7^ (version 1.6.2), and DecontX^3^ (version 1.6.0). All methods were run with default parameters as described in their documentation, except for simulated and negative control datasets.

The methods have different input requirements. Most methods (DecontX, SoupX, FastCAR, and CellClear) by default use both a filtered and an unfiltered count matrix. For droplet-based datasets, the filtered matrix used emptyDrops to detect putative cell-containing droplets. For non-droplets-based datasets the threshold on the number of counts for separating empty droplets from putative cell-containing droplets was manually set. By default, scAR uses only a filtered count matrix, but it has the option to run on unfiltered count matrices. This mode was not included in the benchmark as the run time was unfeasible. By default, CellBender uses only an unfiltered count matrix and automatically detects putative cells. SoupX and DecontX have the option to run using only a filtered count matrix, this mode of operation was included in the benchmark. In addition to count matrices, two methods (scCDC, and SoupX) require clustering information, which was obtained through the standard Seurat v5 pipeline^19^ manually specifying the number of dimensions and clustering resolution (Supplemental Table S6). Subsequently, barcodes with high mitochondrial content were removed.

### Datasets

Seven datasets covering 64 samples and 135.561 cells were used for benchmarking the ambient RNA removal methods (Supplemental Table S1). These datasets span four different dataset categories: species-mixing, strain-mixing, negative control and synthetic. There are three species-mixing datasets, varying in their level of ambient RNA and complexity. The low complexity species-mixing dataset is a one-to-one mixture of fresh frozen human 293T and mouse 3T3 cells. Count matrices were downloaded from 10X Genomics^20^. The medium complexity (GSE147203) samples are mixtures of human islets of Langerhans spiked in with a mix of human Jurkat cells and murine 32D cells^21^. The high complexity (GSE207393) samples are human islets of Langerhans xenografted into murine kidneys^22^. For both the medium and high complexity samples, raw data was aligned to a composite human-mouse genome^23^ and quantified using STARsolo^24^. After quality control, we defined the species of origin by calculating the fraction of UMIs mapped to genes from each species and retained only cells with at least 75% mapped to one species. In these samples, the ground truth for ambient RNA was defined as all UMIs assigned to genes from minor species, and endogenous RNA was defined as all UMIs assigned to genes from the major species.

For the strain-mixing dataset (GSE218853)^25^, which are mixtures of kidney cells from three inbred mice strains from two subspecies (*M. m. domesticus:* C57BL/6 and 129S1/SvImJ and *M. m. castenus*: CAST/EiJ), preprocessed count matrices were downloaded. In these samples, the ground truth for ambient and endogenous RNA was determined as the original authors, with the exception that only genotype-specific single nucleotide polymorphisms (SNPs) with at least 2 counts were included.

The negative control dataset, a smart-seq2 dataset (E-MTAB-5061) from human islets of Langerhans^13^, were aligned to the human genome (2020-A human GRCh38) and quantified using STARsolo^24^. Only three samples (HP1506401, HP1507101, and HP1508501T2D) were able to run through all decontamination runs and therefore selected for further analysis. For these samples, the ground truth for endogenous RNA were defined as all UMIs, as the samples are expected not to contain ambient RNA.

Synthetic datasets were simulated by first fitting gene-wise negative binomial distributions to single-cell RNA-seq data on MCF12A, HCC1500 and HS578T cells^26^. For each real cell to be simulated, we define a total number of UMIs and randomly sampled: a cell type using a specified mixing probability and an ambient fraction for each cell using a normal distribution with a specified average and standard deviation (Supplemental Table S1). Based on the total number of UMIs and ambient fraction, we calculate the number of non-ambient UMIs and sample from the cell type-specific negative binomial distribution. The remaining ambient UMIs were sampled from the two other cell type-specific negative binomial distributions proportionally to the mixing fraction. For each empty cell to be simulated, we define a total number of UMIs and sample from all cell type-specific negative binomial distributions proportionally to the mixing fraction.

### Characterizing ambient RNA

The strain- and species-mixing datasets were used to characterize ambient RNA. For strain-mixing, only features with at least two counts across genotype-specific SNPs were included, as ambient RNA contamination can only be reliably estimated for those. For the medium complexity species-mixing dataset, only the spike-in cells of the opposite species, compared to the source of islets of Langerhans, were included. The major cell types were disregarded, as the spike-ins are low abundant, why there is little-to-no spike-in to islet cell contamination.

For every cell, the fraction of ambient RNA was calculated and associated with regression analyses with cellular parameters, including the number of UMIs, number of features, the fraction of UMIs derived from exons, or from protein-coding, ribosomal, mitochondrial genes. For every feature, we calculated the fraction of ambient RNA and associated it with overall expression levels, expression frequency in cells or empty (less than 50 UMIs) droplets, as well as the fraction of UMIs derived from exons.

### Evaluating ambient RNA removal and endogenous RNA preserved

Decontamination was evaluated by their ability to remove ambient RNA, preserve endogenous signals, and how they affect biological interpretation. An ambient removal score (ARS) and endogenous retain score (ERS) were evaluated by comparing the input to the corrected by each method, calculating their relative differences.

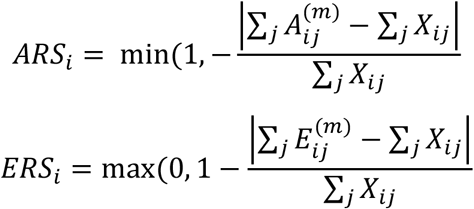

*X*_*ij*_: input ambient (in ARS) or endogenous (in ERS) UMI counts

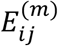: endogenous counts after correction by method m

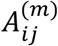: ambient counts after correction by method m

*j*: cell index

*i*: gene index

For the negative control samples and synthetic datasets simulated without ambient RNA, only endogenous retain was calculated, as these datasets do not contain any ambient contamination.

### Evaluating biological interpretation

The impact of ambient RNA removal on biological interpretation was evaluated across several tasks and metrics.

For label transfer, we used Azimuth^15^ and evaluated only islets of Langerhans and kidney-derived cells across datasets, as we had reference atlases available. For species-mixing experiments, we estimated the observed transcriptome with ambient RNA contamination by adding UMIs for genes across species based on orthologs. The label transfer was evaluated by several metrics: the fraction of highly confident (>0.9) mapping and annotation scores, the highest adjusted Rand index between unsupervised clustering (resolutions between 0.05 and 1.0 in steps of 0.05) and transferred labels, the label purity of neighborhoods identified by Milo^27^, the average silhouette width of labels in across the first 20 principal components, as well as the root mean square error (RMSE) of each cell assigned to a label compared the average of all cells assigned to that label for features expressed in at least 10% of the cells in that label.

For sample integration, all datasets were evaluated, except the low complexity species-mixing dataset, as it is only one sample. For each dataset, we integrated the samples using Harmony^28^. The integration was evaluated by several metrics: the highest adjusted Rand index between unsupervised clustering (resolutions between 0.05 and 1.0 in steps of 0.05) and sample labels, the local inverse Simpson index (LISI)^14^, principal component regression and average silhouette width for sample labels in the first 20 principal components, the Gini coefficient of marker gene expression, as well as the Moran I’s and entropy of their expression using nearest neighbors.

For within-sample quality, all datasets were evaluated using several metrics: For species- and strain-mixing datasets, the purity of neighborhoods identified by Milo^27^ in terms of species of origin, as well as the purity of marker genes of neighborhoods, as well as the LISI of species of origin using the first 20 principal components. For all datasets, the Gini coefficient of marker gene expression, as well as the Moran I’s and entropy of their expression using nearest neighbors.

### Ranking methods

To rank methods for each evaluated task (ambient removal, endogenous retain, sample quality, label transfer and batch integration), we averaged across metrics for each dataset and method pair, if more than one metric was used to evaluate the task. Subsequently, the values were min-max scaled across methods for each dataset and task pair and averaged across all datasets within each task. The rank for each task was obtained by ordering the average task scores from highest to lowest. An overall weighted score was obtained by calculating the weighted sum of task score, weighting endogenous retain and ambient removal by a factor of 2, as these tasks are the core functionality of the methods. The overall rank was obtained by ordering the weighted overall scores from highest to lowest. Furthermore, we calculated a score for biological interpretation by taking the median of task scores across the sample quality, label transfer and batch integration tasks. Finally, we calculated an overcorrection score by min-max scaling only control samples based on the difference to the ground truth across all tasks, except ambient removal and averaging across tasks and samples for each method.

## Code availability

The code used to generate the results presented in this study will be made openly available on GitHub at https://github.com/madsen-lab/AmbientRemovalBenchmark. The repository contains all scripts and documentation required to reproduce the analyses.

## Data availability

This study solely used publicly available datasets, all listed in the ‘Methods and Materials section and on the GitHub repository listed in the ‘Code availability’ section.

## Acknowledgements

This work was supported by grants from the Novo Nordisk Foundation (NNF21SA0072102 and NNF21OC0068929), as well as the Danish National Research Foundation (DNRF grant No. 141) to the Center for Functional Genomics and Tissue Plasticity (ATLAS). Computation for this project was performed using the UCloud interactive HPC system, which is managed by the eScience Center at the University of Southern Denmark. We thank all the members of the MadLab group for comments and fruitful discussions that improved the manuscript.

