## Supplemental Tables for "Benchmarking computational decontamination of ambient RNA"

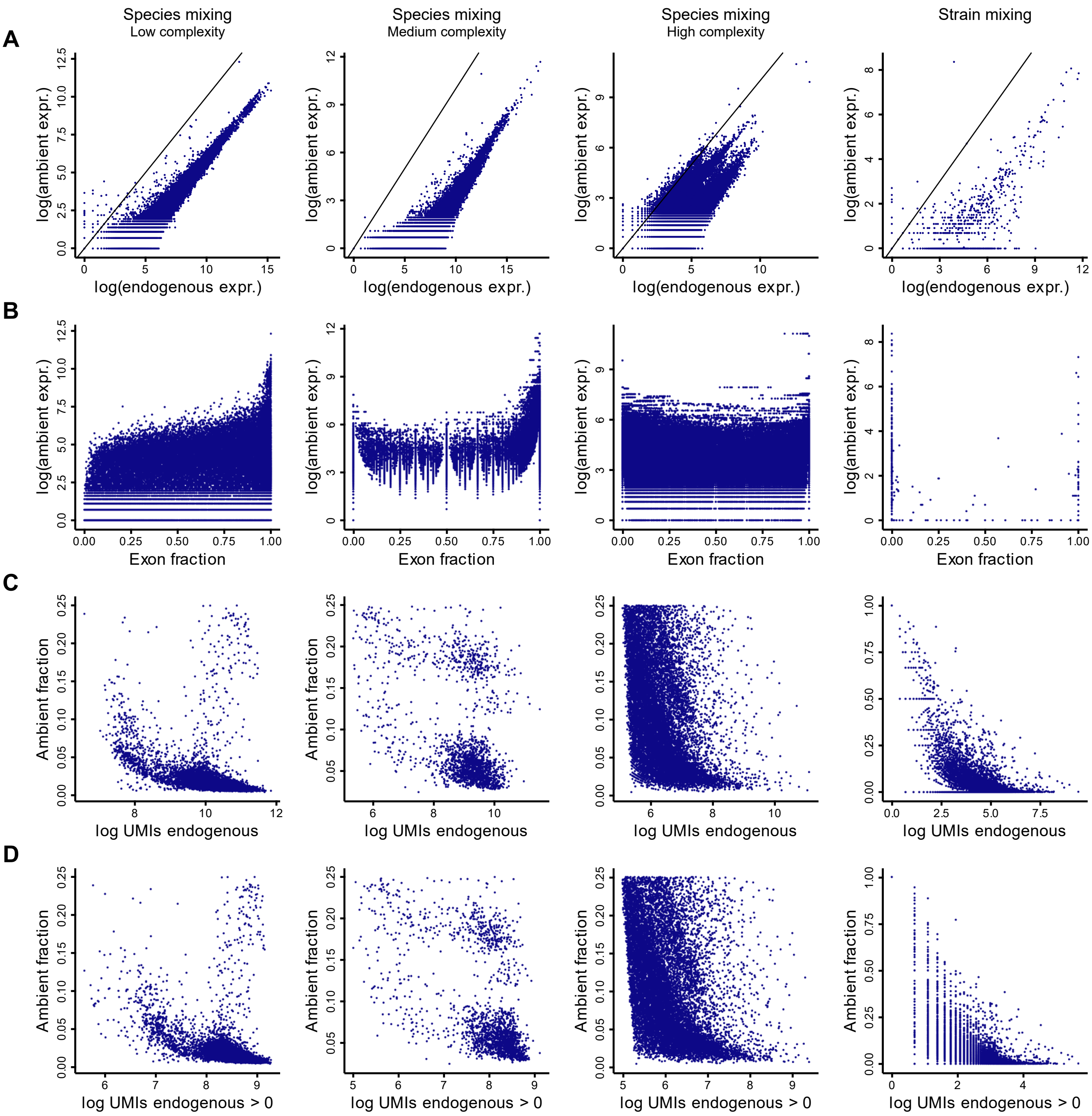


**Supplemental Figure S1 (related to Figure 2). A)** Scatterplot showing the log ambient and endogenous expression level per gene. **B)** Scatterplot showing the log ambient expression level and fraction of UMIs derived from exons per gene. **C)** Scatterplot showing the log total UMI count and the fraction of UMIs derived from ambient RNA per cell. **D)** Scatterplot showing the log total feature count and the fraction of UMIs derived from ambient RNA per cell.

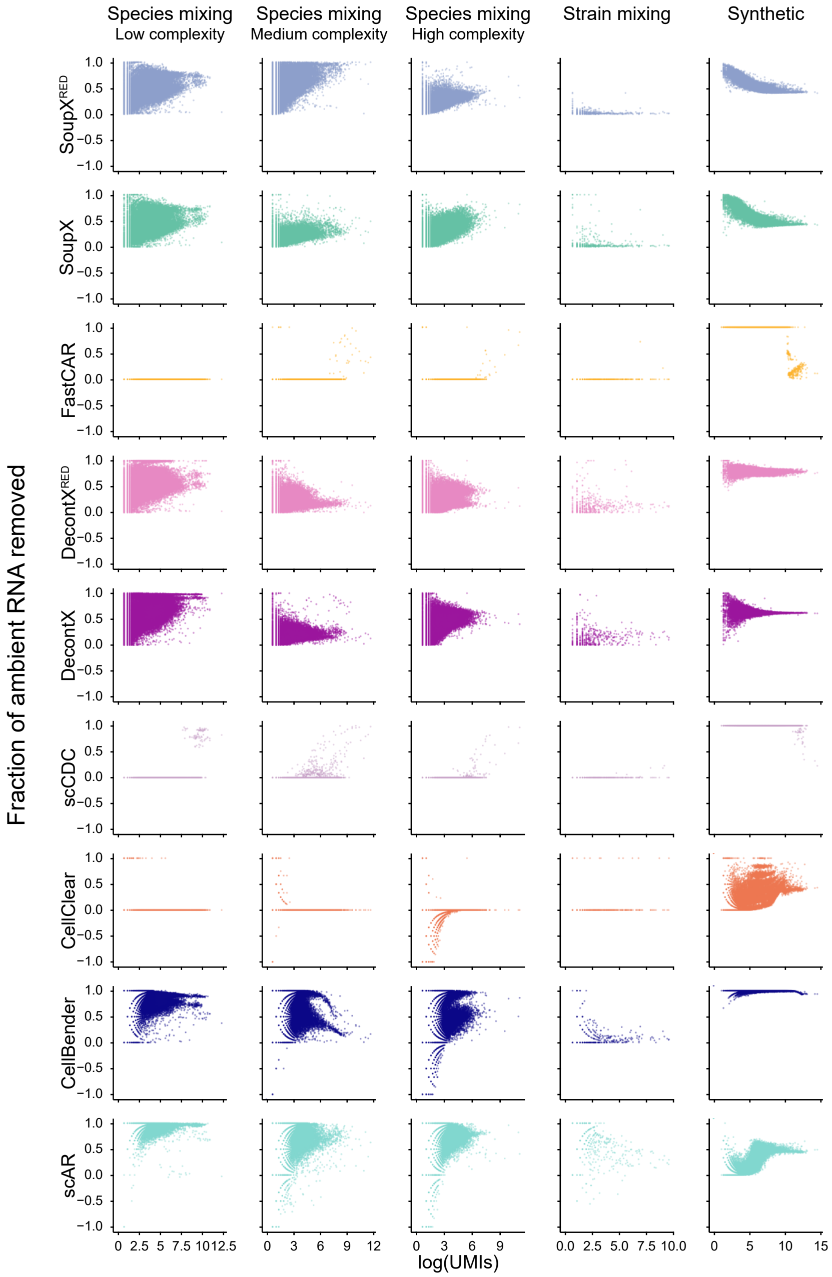


**Supplemental Figure S2 (related to Figure 3).** Scatterplot showing the log endogenous expression and the fraction of ambient RNA removed for the indicated methods in the indicated datasets.

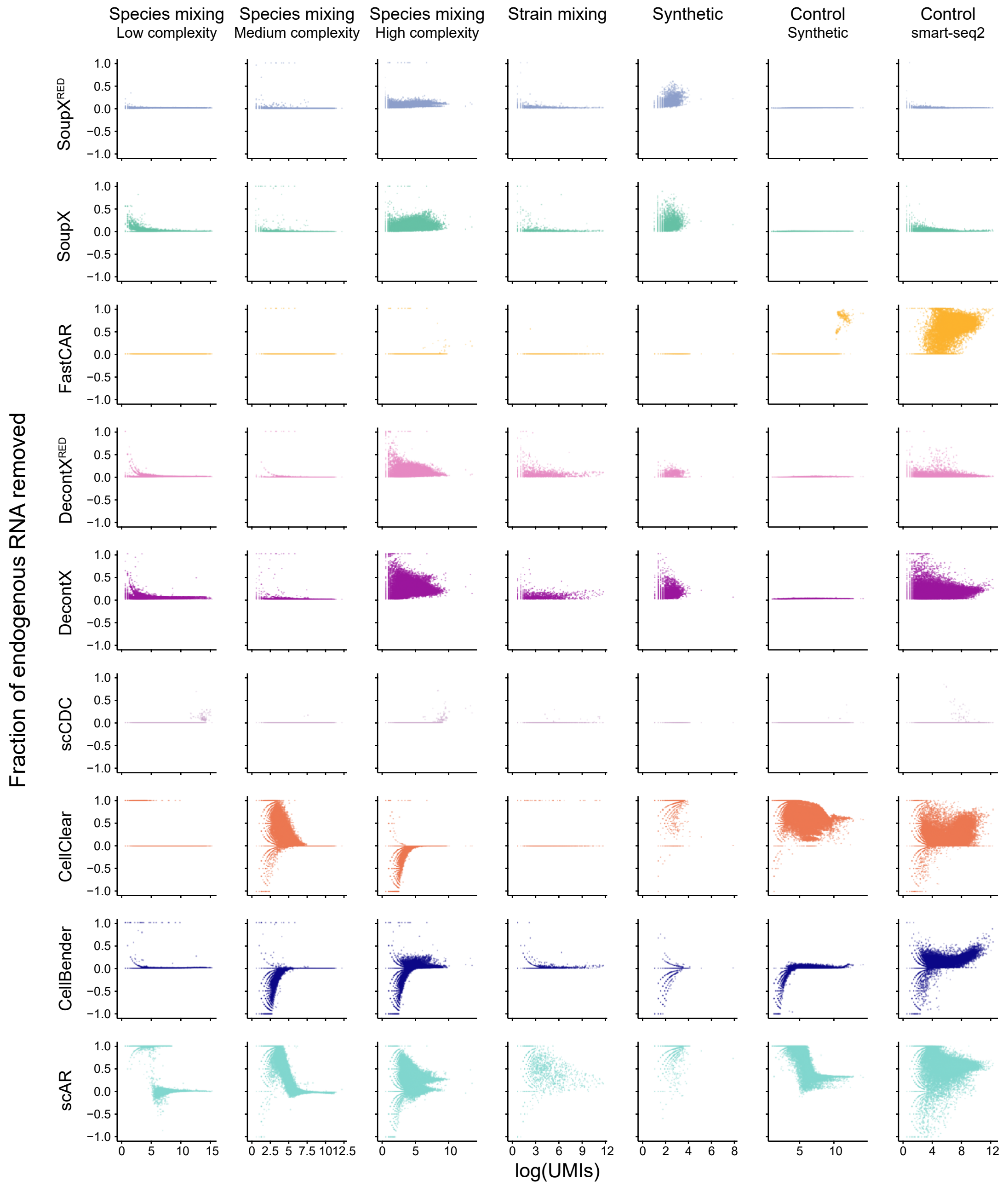


**Supplemental Figure S3 (related to Figure 3).** Scatterplot showing the log endogenous expression and the fraction of endogenous RNA removed for the indicated methods in the indicated datasets.

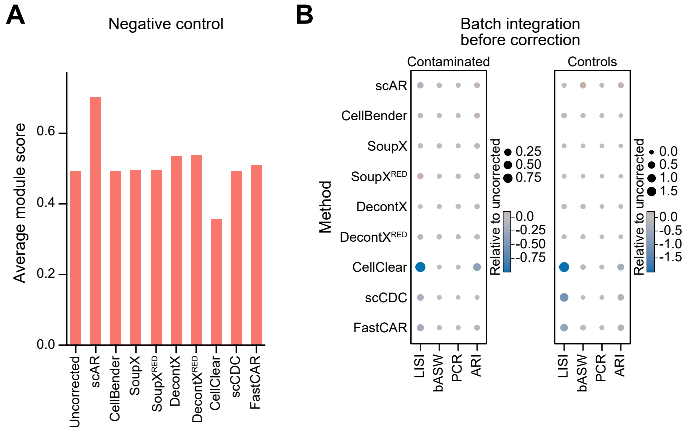


**Supplemental Figure S4 (related to Figure 4). A)** Barplot showing average module scores for marker genes defined in the ground truth dataset from Figure 4A in the same sample without ambient RNA contamination**. B)** Dotplots showing the average of the indicated metrics relative to the uncorrected data for contaminated or control datasets for batch integration prior to integration.
